# NOVA: a novel R-package enabling multi-parameter analysis and visualization of neural activity in MEA recordings

**DOI:** 10.1101/2025.10.01.679841

**Authors:** Caroline C. Escoubas, Ekin Guney, Àlex Tudoras Miravet, Nathan Magee, Ronald Phua, Davide Ruggero, Anna V. Molofsky, William A. Weiss

## Abstract

Multielectrode array (MEA) technology enables simultaneous recording of electrical signals from neuronal networks, producing complex datasets. Current analytical approaches typically examine a limited number of metrics such as mean firing rate and synchronicity, leaving much of the data underutilized. To address this gap, we created NOVA (<u>Neural</u> Output Visualization and Analysis), an accessible R-based computational tool for comprehensive MEA data interpretation and visualization. NOVA integrates dimensionality reduction through principal component analysis, hierarchical clustering with heatmap generation, and temporal trajectory mapping of network activity patterns. Our code offers both a userfriendly pipeline requiring minimal coding background as well as customizable advanced plotting modules for experienced users. Validation experiments using primary cortical neurons during development and pharmacological manipulation demonstrated NOVA’s capacity to detect subtle activity shifts overlooked by conventional methods. Notably, our unbiased approach identified network burst duration as a stronger contributor to activity variance than commonly reported firing rate metrics, exemplifying NOVA’s utility for discovering meaningful patterns and generating data-driven hypotheses.

## Introduction

MEA (multielectrode arrays) technology measures voltage changes in the extracellular environment of electrically active cells ^1–3^. The large area covered by electrodes in multi-well MEA records simultaneously from thousands of neurons, suitable for analyzing network activity and for high-throughput screens^1^. MEA characterize the dynamics of electrical activity in in the context of neurodevelopment^3,4^, neuroscience^5–7^, toxicology^8^ neurodegeneration^9,10^ and neuro-oncology^11,12^. MEA electrophysiology recordings produce high-dimensional datasets that challenge conventional analytic approaches. While over 30 outputs are measured during MEA recordings, most studies focus on only mean firing rate and synchronicity, failing to realize the full potential of the system, and potentially missing critical information regarding the regulation of neuronal activity.

Here we developed a user-friendly analytic pipeline in R, called NOVA (<u>N</u>eural <u>O</u>utput <u>V</u>isualization and <u>A</u>nalysis), which provides a standardized statistically rigorous computational framework for transforming batched MEA recordings into interpretable visualizations. NOVA generates easily interpretable graphs that reflect changes in the dynamics of neuronal activity and is useable by scientists without a background in coding. We provide a detailed explanation of functions embedded into NOVA. Applying NOVA to MEA analysis of primary neuronal cultures, we highlight NOVA’s ability to capture changes in neuronal activity over time revealing patterns of activity changes.

### The NOVA package effectively captures changes in neuronal activity dynamics

NOVA contains functions which can normalize the data exported from the Axis Navigator Software. Using the individual metrics measured in MEA (**Supp Table 1**), NOVA performs a principal component analysis (PCA) with dimensionality reduction, and statistical computations, plots heatmaps with multiple scaling options and hierarchical clustering and calculates trajectory analysis representing the evolution of neuronal activity through principal component space. A detailed explanation of each function used to generate these graphs is provided in the Methods section. We developed a “main” version of the code which can be run without any prior R coding experience and will generate a number of default graphs to visualize differences in patterns of neural activity. We also provide highly flexible and adjustable “Enhanced” plotting modules for Heatmaps, Trajectories and PCA plots for more experienced R users. To validate our NOVA package, we analyzed MEA recordings from primary cortical neuron cultures either during maturation or after treatment with neuronal agonists and antagonists.

#### Neuronal activity agonists distinguish overall changes in patterns of neuronal activity

Primary neuron cultures were prepared from postnatal day 0-2 C57Bl6 pups (see *Methods*), plated on MEA plates and matured for 12 days (days in vitro; DIV12), a timepoint where the culture is morphologically and electrophysiologically connected and mature^3,4,13^ (**Fig1A**). Cultures were then treated, and neuronal activity recorded using the Axion Biosystems platform at several points during the first 2h post treatment to capture acute dynamic responses to drugs. We treated these WT neuron cultures with neuronal activity agonists that target the major classes of synaptic receptors: NMDA 20μM (NMDA receptor agonist)^14^, AMPA 10μM (AMPA receptor agonist), kainic acid 1μM (kainite receptor agonist)^15^, DHPG 10μM (metabotropic receptor agonist)^16,17^ and gabazine 5μM (GABA receptor antagonist)^18,19^ (**Fig1B**). We optimized the concentration for each of drug to increase neuronal activity without inducing excitotoxicity. Raster plots generated at 2h post-treatment showed strong qualitative differences between the neuronal activity patterns induced by different drugs.

To capture the complexity and multivariate nature of the MEA data, rather than analyze isolated outputs, we performed PCA analysis, including all the drug treatments and all the timepoints (**Fig 1 C,D, Supp Fig1 A,B**). This 2-dimensional representation highlights separation between treatments with the most differences observed in the DHPG, gabazine and NMDA treated cultures. Notably, among the top 20 variables contributing to PCA variance (**SuppFig1A**), mean firing rate and weighted mean firing rate ranked 15^th^ and 16^th^ respectively, consistent with published work^4^, suggesting that selectively looking at these two metrics would fail to capture the complexity of neuronal responses to stimuli. Heatmaps generated with the NOVA package allow for a more in-depth analysis of the various metrics contributing to the drug effects (**Fig1F**) and show correlated metrics (**SuppFig1C**). Finally, to capture the temporal dynamics of the neuronal responses to drug, NOVA calculates and generates trajectory plots (**Fig 1G**). Interestingly, our data illustrates that most of the effect is seen during the first 30min in DHPG treated neurons. Unlike DHPG, gabazine and NMDA treated neurons continue to change their pattern of activity until T=2h, particularly along principal component 1 which has high contributions from network burst frequency and synchrony index metrics (**SuppFig1B**).

**Figure 1:**
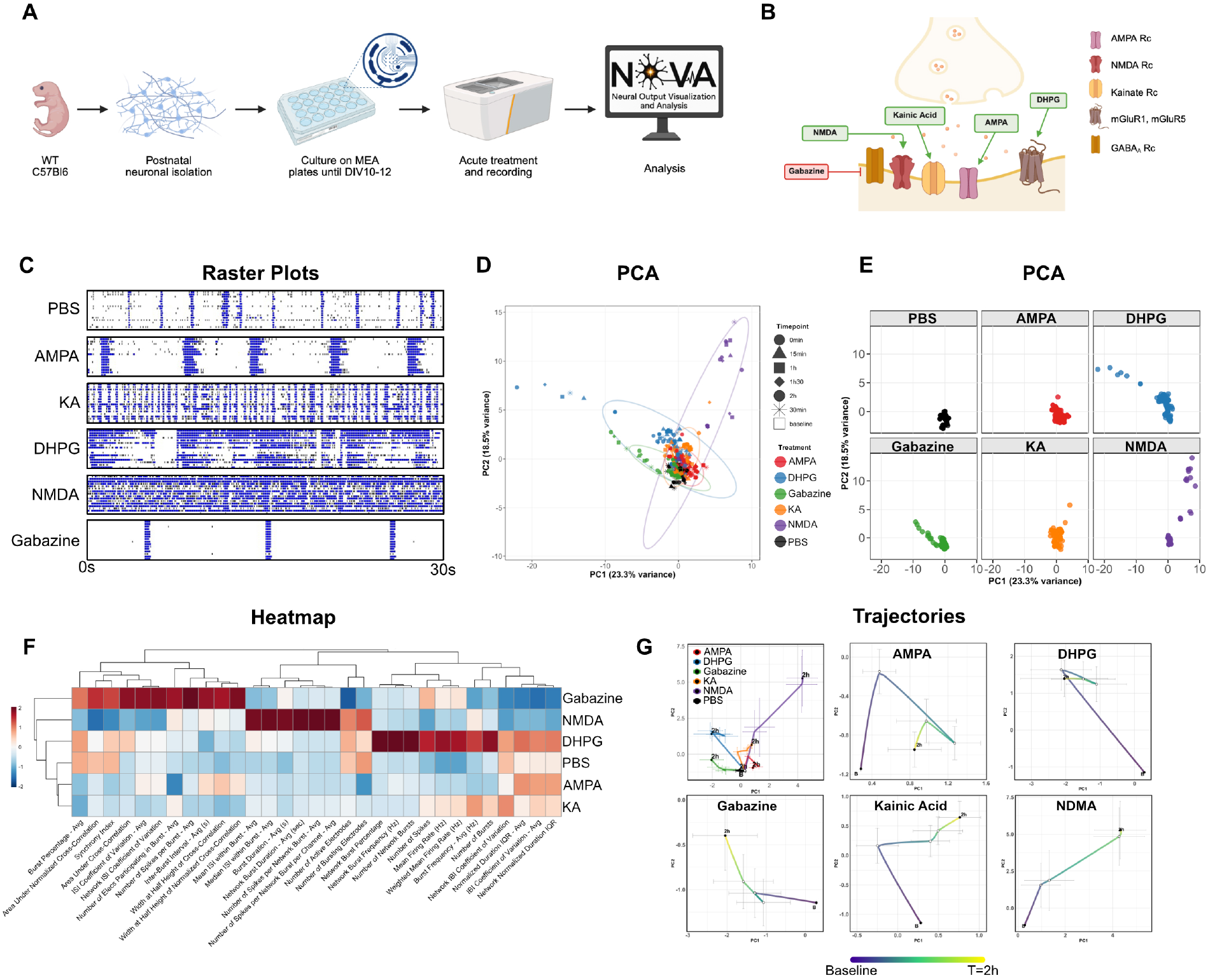
Analysis of MEA recordings after induction of neuronal activity using NOVA. **(A)** Experimental design. Recordings were obtained at baseline, and 0min, 15min, 30min, 1h, 1 h30min and 2h after drug treatment. **(B)** Schematics of the neuronal activity agonists used and their primary synaptic target: gabazine (GABA Rc antagonist), NMDA (NMDA Rc agonist), kainic acid (Kainate Rc agonist), AMPA (AMPA Rc agonist), DHPG (mGluR1, mGluR5 agonist) **(C)** Representative raster plots for each treatment used (30s duration). Each horizontal line represents one electrode (16 total). Blue shows bursts. Black shows individual spikes. **(D)** Principal component analysis on all data combining treatments (different colors) and timepoints (different shapes) with 95% confidence ellipses using NOVA. Each dot represents a well. **(E)** PCA plots split by treatment using NOVA. Each dot represents a well. **(F)** NOVA heatmap with hierarchical clustering with combined timepoints after drug treatment. **(G)** NOVA analysis of average trajectories representing responses over the first 2h post-treatment.

Our novel R-package NOVA allows users to distinguish overall changes in patterns of neuronal activity using principal component analysis. Heatmaps and trajectory analysis then provide data to choose specific metrics and timepoints (**SuppFig1D-G**) to generate precise hypothesis and potentially perform mechanistic follow-up studies.

#### Neuronal activity antagonists illustrate the ability of NOVA to analyze conditions with different effect sizes

We next treated WT neuronal cultures with neuronal activity antagonists to further validate our novel pipeline. We chose antagonists that target major classes of synaptic receptors: NBQX 10μM (AMPA receptor blocker), ACET 10μM (kainite receptor blocker), D-AP5 10μM (NMDA receptor blocker) and GABA 5μM (GABA receptor agonist) (**Fig 1A**). Similarly to the agonist drugs, we optimized the concentration of each drug to achieve a reduction in neuronal activity without complete ablation of neuronal firing (**Fig2B**). Representative raster plots isolated from 2h timepoint illustrate the overall decrease in spiking activity and change in network burst profile (**Fig 2B**). Principal component analysis combining all treatments and timepoints distinctively separates GABA and NBQX treated groups with less separation in the PCA plot between ACET and D-AP5 treated groups compared to PBS control (**Fig2C, SuppFig2A,B**). The combined timepoints heatmap highlights a significant decrease in network burst frequency in GABA and NBQX treated neurons, consistent with the raster plot visualization (**Fig2D, SuppFig2C**). Overall, the ACET and D-AP5 treated neurons do not show large differences compared to the PBS group in the heatmap, rather suggesting dampened neuronal activity without a complete change in activity pattern (**Fig2D**). Trajectory plots, while confirming the distinct effect of NBQX and GABA on neurons, suggest that ACET and D-AP5 induce acute changes before reverting to a position close to the PBS control in this 2-dimensional representation (**Fig2E**). Combining the information from the heatmap and the trajectory plots, we looked at the dynamics of a single metric, mean firing rate over time. Indeed, we observed that mean firing rate was acutely decreased at time points 0min, 15min and 30min (**SuppFig2E**) before going back to baseline (**SuppFig2D,E**). These data illustrate the applicability of our pipeline to analyze conditions with different effect sizes.

**Figure 2:**
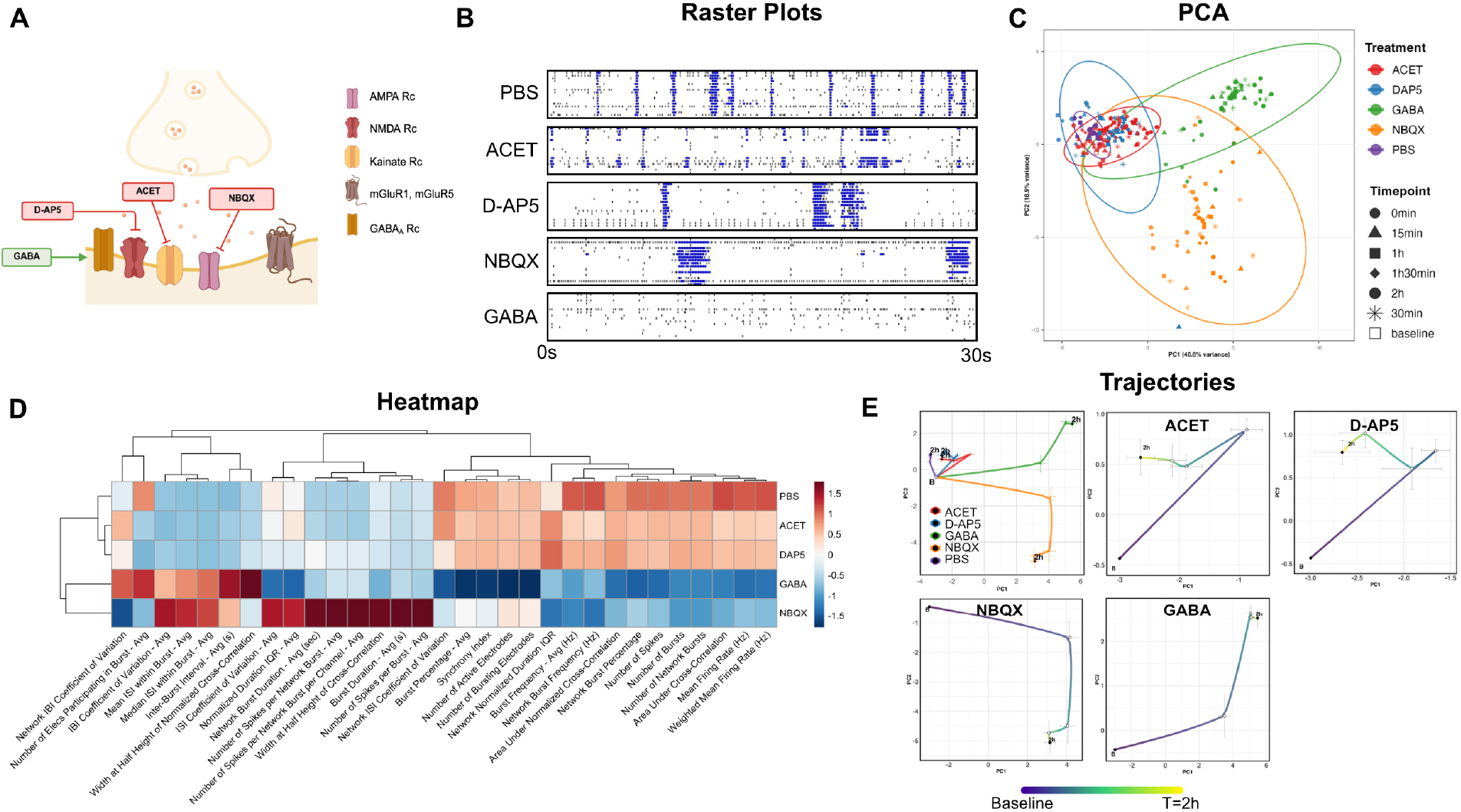
NOVA analysis of MEA recordings after repression of neuronal activity. **(A)** Schematics of neuronal activity agonists and their primary synaptic targets: GABA (GABA Rc agonist), D-AP5 (NMDA Rc antagonist), ACET (Kainate Rc antagonist), NBQX (AMPA Rc antagonist). **(B)** Representative raster plots for each treatment (30s duration). Each horizontal line represents one electrode (16 total). Blue shows bursts. Black shows individual spikes. **(C)** Principal component analysis on all data combining treatments (different colors) and timepoints (different shapes) with 95% confidence ellipses using NOVA. Each dot represents a well. **(D)** NOVA heatmap with hierarchical clustering at combined timepoints after drug treatment. **(E)** NOVA analysis of average trajectories representing responses over the first 2h post-treatment.

#### Analysis with NOVA reveals distinct stages of primary cortical neuron culture maturation

Beside comparing treatment effects and kinetics, another common use of MEA analyses is to study the rate of neuronal functional maturation *in vitro* between genetic backgrounds^4^. Unlike analysis of treatment effects for which we normalize data to the baseline value in each well (**Fig1, Fig2**), comparing maturation rates requires analysis of raw MEA data. To validate our pipeline, we recorded neuronal activity every day from DIV2 to DIV13, in MEA plates seeded with WT neurons (**Fig3A**). Morphological maturation of neurons in culture (**Fig3B**) correlated with gradual increase in number of spikes and emergence of network burst firing (**Fig3C**), as has been extensively characterized^3,4,13^. Principal component and trajectory analyses revealed a distinct temporal separation of the data along principal component 1 (**Fig3D-F**) with major contributions from variables related to network burst firing (**Fig3E**). The heatmap representation of these data revealed an interesting pattern of maturation. Rather than a completely linear progression over time, patterns of neuronal activity clustered into 3 groups: DIV2-5 where individual, unsynchronized spikes were predominant, DIV6-11 where synchronized firing and network burst emerged and DIV12-13 where bursts were longer and the activity was more synchronized, likely reflecting a more connected and mature culture (**Fig3G**).

**Figure 3:**
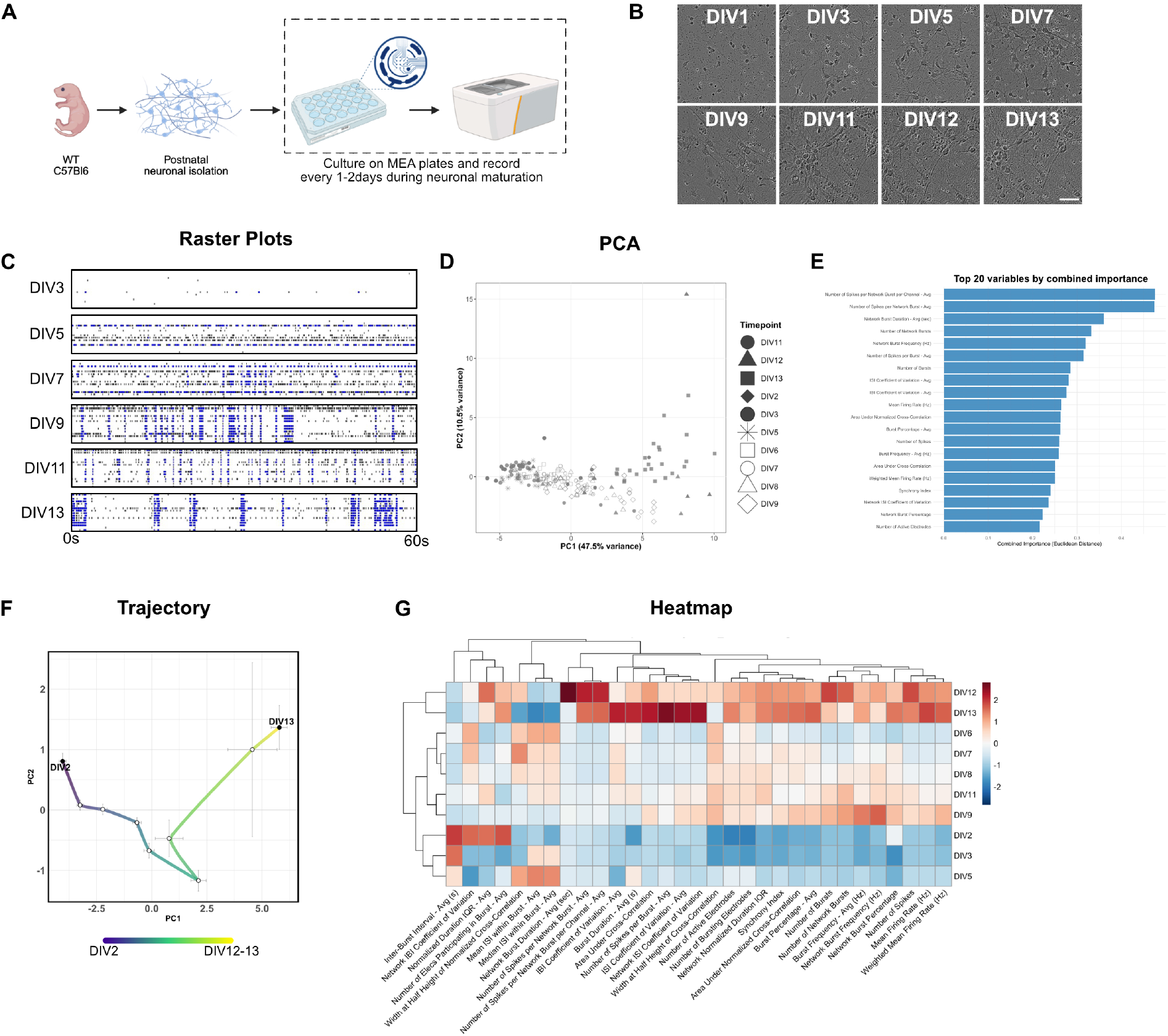
Analysis of MEA recording during cortical neuron maturation *in vitro* using NOVA. **(A)** Schematics of the experimental design. **(B)** Representative brightfield images of primary cortical neurons at timepoints DIV1, 3, 7, 9, 11, 12, 13, showing neuronal maturation and formation of processes (scale bar = 50μm). **(C)** Representative raster plots for selected timepoints during maturation (60s duration). Each horizontal line represents 1 electrode (16 total). Blue shows bursts. Black shows individual spikes. **(D)** NOVA representation of principal component analysis on all data combining timepoints (different shapes) with 95% confidence ellipses. Each dot represents a well. **(E)** NOVA representation of relative contributions of top 20 variables to Principal Component 1 and Principal Component 2 for the neuronal maturation PCA analysis. **(F)** NOVA average trajectory representing cortical maturation over 13 days *in vitro*. **(G)** NOVA heatmap with hierarchical clustering at different timepoints during cortical maturation *in vitro*.

## Discussion

Development of MEA technology and in particular multi-well MEA plates represent important tools to assess network level activity, particularly in cultured neurons, enabling high throughput experiments. Because MEA recordings generate a large number of metrics, advanced analytic methods are important to broaden applicability. Others in the field have published statistical methods and code to either dissect indepth raw MEA recordings, run lower dimensional analysis such as principal component analysis or develop index scores^4^. While valuable, analysis of individual waveforms from raw MEA recordings can be difficult to implement and interpret for non-electrophysiologists. Alternatively, aggregating neuronal activity metrics into composite scores, while providing high level overview, will often fail to capture fine scale variations. Here we developed a streamlined and easy to use code, NOVA, to provide comprehensive analysis and plotting options for MEA recordings. NOVA generates plots such as PCA, heatmaps and trajectories allowing for visually impactful and easy to understand representations of changes in patterns of neuronal network activity. These analyses will effectively highlight broad differences between groups while providing the option to analyze the relative contribution of individual metrics, such as burst frequency or number of spikes. In our neuron cultures treated with synaptic agonists, the analysis of the relative contribution of different metrics to principal components 1 and 2 in our PCA analysis reveals the low contribution of mean firing rate (ranked 15^th^) and the higher contribution of network burst duration (ranked 3^rd^). Over relying on mean firing rate as often seen in the literature would therefore fail to capture the dramatic effect seen in NMDA treatment for example. Our pipeline can therefore be used to analyze changes in patterns of activity in an unbiased way and identify which feature of the network activity is most altered and use those observations for hypothesis generation.

The *in vitro* validation of our NOVA package focused on optimized concentrations of both neuronal activity agonist and antagonist drugs. We chose concentrations that functionally modulated network activity without inducing excitotoxicity and complete ablation of action potentials. Notably, the concentrations of agonists used in this manuscript are generally lower than what can be seen in the literature, where validation of these drugs often rely on immediate early genes such as *cfos*, staining. Our data therefore provides a useful resource to modulate neuronal activity in primary cultures.

Finally, recent evidence points to the role of non-canonical neuronal pathways, such as immune pathways, in regulating brain development, behavior and neuronal activity^20–22^. The ease of MEA recordings paired with our novel NOVA R-package should allow non electrophysiologists to venture into neural network biology. Our principal component analysis, trajectory analysis or heatmaps can easily identify changes in patterns of activity, useful when studying effect of cytokines on neuronal cultures or the impact of modulation of immune genes in monoculture or co-culture settings.

## Materials and Methods

### Data and code availability

NOVA R-package code used to analyze MEA recordings datasets is available on GitHub (github.com/atudoras/nova) and can be loaded in R via library(NOVA) after installation. Any additional information required to reanalyze the data reported in this paper is available from the corresponding authors upon request.

### Animals

All mouse strains were maintained in the University of California, San Francisco specific pathogen–free animal facility, and all animal protocols were approved by and in accordance with the guidelines established by the Institutional Animal Care and Use Committee and Laboratory Animal Resource Center. Mice were housed in a 12-hour light/dark cycle (7am-7pm) at 68-79°F and 30-70% humidity. Male and female mice were group-housed when possible, with up to 5 mice per cage. Littermate controls were used for all experiments and both male and female pups were used for the neuronal isolations. The following mouse strains used are described in the table below and are referenced in the text: C57BL/6J (Jax#000664).

### Primary neuronal cultures

Primary cortical neural cultures were established from brains of C57BL/6J mice pups (P0-P2) in accordance with UCSF Institutional Animal Care and Use Committee (IACUC). Cortices were dissected and collected in serum-free neural conditioned media (NCM, see recipe below). Papain, ovomucoid and DNAse are prepared according to manual instructions of Worthington Papain Dissociation System (#LK003153). NCM was completely removed from dishes with mouse cortices and 1mL papain-DNAse mix was added. Cortices were minced to 1-2 mm^2^ pieces with a blade and transferred to a 15ml falcon tube. The remaining Papain-DNAse mix was added to the falcon tube and incubated at 37°C for 20min, with gentle swirling of the tube every 3min. Mechanical dissociation was performed by trituration using sterile P1000 pipette tips with 3 different size cut tips. Neural cell suspension was centrifuged at 200g for 5min at 4°C. The supernatant was discarded, and the pellet was resuspended in resuspension buffer (Worthington Papain Dissociation System) and filtered through 40um cell strainer. The filtered neural cell suspension was slowly added on top of the blocking buffer (Worthington Papain Dissociation System) to create a gradient and centrifuged at 70g for 6min at RT. Following centrifugation, the supernatant was discarded, and the pellet was resuspended in NCM. Cell counts and cell viability was assessed using a Bio-Rad TC-20 automatic cell counter and neurons were resuspended at a concentration of 10 x 10^6^/mL. Droplets of 10ul with 100,000 viable neural cells in NCM were plated at the center of the electrode area according to Axion BioSystems MEA plating guidelines with 7ml 1xDPBS added between the wells and incubated at 37°C for 1 hour to allow cell adhesion. After 1 hour, 400ul of NCM was added to each well. The entire 400ul NCM media was replaced at DIV1 and half of the media (200ul) was replaced at DIV4, 8 and 11/12 until neurons reached functional and morphologic maturity and connectivity.

#### NCM

Neurobasal-A (Gibco #10888022), B-27™ Supplement 1X (Gibco #17504001), Antibiotic-Antimycotic 1X (Gibco #15240096), CTS GlutaMAX-I Supplement 1X (Gibco #A1286001), Sodium Pyruvate 100 mM (Gibco #11360070) and Mycoplasma Elimination Reagent 1:20,000 (Plasmocure #ant-pc).

### Multi Electrode Array plate preparation

MEA plates (CytoView 24-well plates with 16 electrodes, Axion BioSystems #M384-tMEA-24W) were coated with PDL (20 ug/ml, ThermoFisher Scientific #A3890401) at RT overnight, washed 3x with 1x DPBS (Gibco #14190144) and coated with laminin mouse protein (2.4 ug/ml, ThermoFisher Scientific #23017015) and Fibronectin Human Protein (0.4 ug/ml, ThermoFisher Scientific #33016015) for 2h at 37C on the day of the neuronal isolation. Following coating, plates were washed 1x with DPBS and air dried in the tissue culture hood with lids open for 30-45min, until plating of the neurons.

### Multi Electrode Array recordings

Neural field potential recordings were performed using the Maestro Edge MEA platform (Axion BioSystems, USA) at 37°C and 5% CO2. Real-time data was collected simultaneously across all 384 electrodes using Neural Real-Time spontaneous activity module in Axion Integrated Studio (AxIS) Navigator software (16 electrodes per well, 24 well plate, 12.5 kHz sampling rate, 200-3000 Hz band-pass). Analysis was performed using AxIS Navigator software (Axion BioSystems). The neuronal events were defined by applying an adaptive threshold crossing method (5 standard deviations of the noise level threshold for spike detection), which defines spike as activity exceeding this threshold. All analysis was performed on only active channels, (exhibited ≥5 spikes per min). The NeuroPlot Tool was used to extract representative raster plots.

### Neuron cultures drug treatment

Neuronal cultures were treated between DIV12-DIV15. After letting the plate equilibrate for 15min in the Maestro Edge MEA platform, one 10min baseline measure was recorded. Plates were then briefly taken out of the Maestro Edge MEA platform (<30sec) and drugs were added in a 10ul volume/well to achieve the listed concentrations. After a brief 1min equilibration, measures (5min each) were taken at T=0min, 15min, 30min, 1h, 1h30 and 2h post drug treatment.

In this manuscript, we optimization the concentrations of the following drugs:

| Abbreviation | Compound name | Concentration | Catalog # |
| --- | --- | --- | --- |
| KA | Kainic acid | 1μM | Tocris/Cat#0222 |
| (S)-3,5-DHPG | 3,5-dihydroxyphenylglycine | 10μM | Tocris/Cat#0805 |
| NMDA | N-methyl-D-aspartate | 20μM | Tocris/Cat#0114 |
| (S)-AMPA | α-amino-3-hydroxy-5-methyl-4-isoxazolepropionic acid | 10μM | Tocris/Cat#0254 |
| Gabazine | SR 95531 | 5μM | Tocris/Cat#1262 |
| ACET |  | 10μM | Tocris/Cat#2728 |
| D-AP5 | D-2-Amino-5-Phosphonovaleric Acid | 10μM | Tocris/Cat#0106 |
| NBQX disodium salt | 2,3-Dioxo-6-nitro-1,2,3,4-tetrahydrobenzo[f]quinoxaline-7-sulfonamide | 10μM | Tocris/Cat#1044 |
| GABA | gamma-aminobutyric acid | 20μM | Tocris/Cat#0344 |

All drugs were resuspended according to manufacturer’s guidelines and were then diluted in PBS to use in MEA recordings. All resuspended drugs, except Kainic acid (4°C), were kept in aliquots at -20°C.

### Neuron cultures developmental analysis

For developmental analysis, we recorded the MEA plates plated with primary cortical neurons starting at DIV2-3 and recorded them every 1-2 days until DIV12-13. For each recording, the plate was left to equilibrate 15min at 37°C and 5% CO_2_ in the Maestro Edge platform before a 10min measure was recorded. On the days of scheduled media change, recordings were made prior to the media change to avoid measuring a transient burst in activity due to addition of fresh media.

### Data processing

MEA recordings were processed using the AxIS Navigator software (Axion BioSystems) and electrode spike features were exported for each timepoint in.csv format and used as input data for NOVA R-package functions. Rows “Genotype” and “Exclude/Include” were manually added to the.csv file. For each well, genotype, treatment and exclusion (if necessary) were manually added. MEA.csv files were renamed to contain the experiment ID and the timepoint of recording.

### NOVA

#### Software development and implementation

NOVA R-package was created using RStudio version (2025.05.1+513). NOVA can be installed from the following GitHub repository: github.com/atudoras/nova. A detailed explanation for each function embedded into NOVA is available below. Example workflows and instructions on how to apply NOVA R-package are available on GitHub at github.com/atudoras/nova. We utilized Claude AI (Anthropic, version Claude Sonnet 4.5) to assist with debugging and troubleshooting technical errors in R code development.

NOVA R-package will be maintained and feedback provided via email or via the GitHub issues tab will be considered to update the code.

#### NOVA Data Processing and Quality Control

Analysis of raw MEA recordings using the Axion Biosystems software generates standardized CSV files containing measurements and individual well identifiers. Users manually add experimental variables (e.g., treatment, genotype, condition, etc) and well as exclusion flags into designated rows. Example data files with the expected structure are provided in GitHub (github.com/atudoras/nova). NOVA implements automated data discovery through (**discover_mea_structure**), which scans experimental directories to extract timepoint information and experiment ID and extracts all relevant metadata from the CSV files. The (**process_mea_flexible**) function implements a core data integration pipeline, reading all selected CSV files and extracting both neural measurements and user-defined experimental metadata from their respective positions within each file. The system automatically applies quality control measures, excluding flagged wells and filtering out standard deviation measurements while preserving mean values to reduce data dimensionality. When specified, the system performs baseline normalization by calculating fold-change values relative to a designated reference timepoint. The system consolidates equivalent timepoint labels and maintains complete audit trails for reproducibility. Quality control operates at multiple stages, applying variance-based filtering to remove low information variables and implementing completeness thresholds to ensure sufficient data. Final datasets are exported with separate sheets for raw and normalized data, ensuring reproducibility and facilitating downstream statistical analysis across all user-defined experimental conditions.

#### NOVA Principal Component Analysis and Dimensionality Reduction

The codebase implements a two-stage PCA pipeline comprising analytical and visualization components. The pca_analysis_enhanced function handles data preprocessing, dimensionality reduction, and statistical computations, while pca_plots_enhanced generates publication-ready visualizations from the analytical results.

The analytical component (**pca_analysis_enhanced**) extends R’s function (**prcomp**) with MEA-specific data handling capabilities. It accepts multiple input formats including processing results, Excel files, or direct data matrices, automatically detecting and validating data structure requirements. The implementation applies singular value decomposition to preprocessed data matrices, offering configurable scaling options and multiple missing value strategies including mean imputation, median imputation, or complete-case analysis. Variance explained calculations quantify the proportion of total variance captured by each principal component, with automated component selection based on cumulative variance thresholds.

The visualization component (**pca_plots_enhanced**) transforms PCA results into multiple plot variants optimized for scientific publication. It generates up to five distinct visualizations including primary and secondary aesthetic combinations, simplified color-only displays, statistical overlays with 95% confidence ellipses computed from group means and covariance matrices in principal component space, and faceted displays for complex experimental designs. The plotting system incorporates colorblind-accessible palettes and supports selective grayscale highlighting of control conditions for enhanced visual contrast.

#### NOVA Temporal Trajectory Analysis

The (**plot_pca_trajectories_general**) function, a key methodological advancement, visualizes neural activities as continuous trajectories through principal component space rather than as static snapshots. Individual wells are identified using pattern matching, tracking the evolution of activity across timepoints with modulable temporal ordering that recognizes standard biological nomenclature.

Individual trajectories receive optional smoothing through a hybrid approach combining linear interpolation with spline fitting (75%/25% weighting) to reduce noise while preserving temporal dynamics. Group-averaged trajectories incorporate standard error calculations across individual wells, enabling statistical assessment of treatment effects. The function generates multiple complementary graphs: individual trajectory plots revealing within-group heterogeneity, group averages with statistical uncertainty, and combined plots for direct comparison across experimental conditions.

### NOVA Heatmap Analysis and Data Aggregation

NOVA’s heatmap functionality through (**create_mea_heatmaps_enhanced**) provides additional insights via the (**pheatmap**) framework. This analysis aggregates measurements across all timepoints within each experimental condition, generating static summaries that highlight persistent treatment differences. This temporal consolidation contrasts with trajectory analysis, which preserves time-course information. Multiple scaling approaches include: z-score normalization for relative comparisons, min-max scaling for absolute value analysis, and robust scaling using median absolute deviation for outlier resistance. Hierarchical clustering employs Euclidean distance, correlation distance, or Manhattan distance metrics with automatic dendrogram generation.

### NOVA Variable Importance Assessment

The (**analyze_pca_variable_importance_general**) function quantifies individual parameter contributions to experimental effects through comprehensive loading-based metrics. The system extracts loadings from the (**prcomp**) rotation matrix, calculates combined importance scores using Euclidean distance in loading space, and determines contribution percentages through normalized sum-of-squares analyses. Multiple visualization options include: loading biplots display variables in principal component space with size-encoded importance, ranked bar charts highlight top contributors, comparison plots show loading magnitudes across components, and heatmaps provide color-coded loading matrices. This functionality enables researchers to identify specific electrophysiological measurements that drive observed group differences.

#### NOVA Implementation and Reproducibility

NOVA maintains methodological consistency through comprehensive parameter logging, automated export of analysis metadata, and standardized output formats. The modular architecture supports workflow customization while preserving analytical rigor. Automated plot generation enables batch processing with configurable resolution and dimensions, while robust error handling ensures stable operation across computational environments. Quality assurance mechanisms validate data integrity at each processing stage, from initial file parsing through final visualization. The system provides detailed progress reporting and maintains complete provenance tracking, enabling full reproduction of analytical workflows. This integration of statistical rigor with practical accessibility makes advanced MEA analysis tractable for neuroscience researchers while meeting computational standards required for rigorous data science applications.

#### NOVA Fully customizable plotting modules

For more experienced users, we additionally created enhanced plotting modules (PCAPlot_module, PCAImportance_module, Trajectories_module and Heatmap_module) which are available in the package’s GitHub repository with usage examples (github.com/atudoras/nova). These modules allow users to entirely customize the data being displayed and the aesthetics of each plot.

#### Statistical analysis

Graphpad Prism 9.3.1 was used for all statistical analyses of individual metrics. Statistical tests used, number of n replicates and p values are described in figures and figure legends. All statistical analyses unless otherwise noted were performed on means of multiple technical replicates (well) per experiment. The NOVA R-package was used for statistical analysis of MEA data.

## Author contributions

Conceptualization: C.C.E., E.G.; Methodology: C.C.E., E.G. and A.T.M.; Investigation: C.C.E., E.G. and A.T.M.; Technical help with dissections: R.P. and N.M.; Writing – original draft: C.C.E., E.G. and A.T.M.; Writing – review & editing, all co-authors; Funding acquisition: W.A.W., A.V.M., D.R.; Resources: W.A.W.; Supervision: C.C.E and W.A.W.

## Acknowledgments

We thank the Pollen lab (UCSF) for access to their Mastro Edge MEA platform. We are grateful for the Weiss, Molofsky and Seeley lab for helpful comments and feedback.

---

**Supplemental Figure 1:**
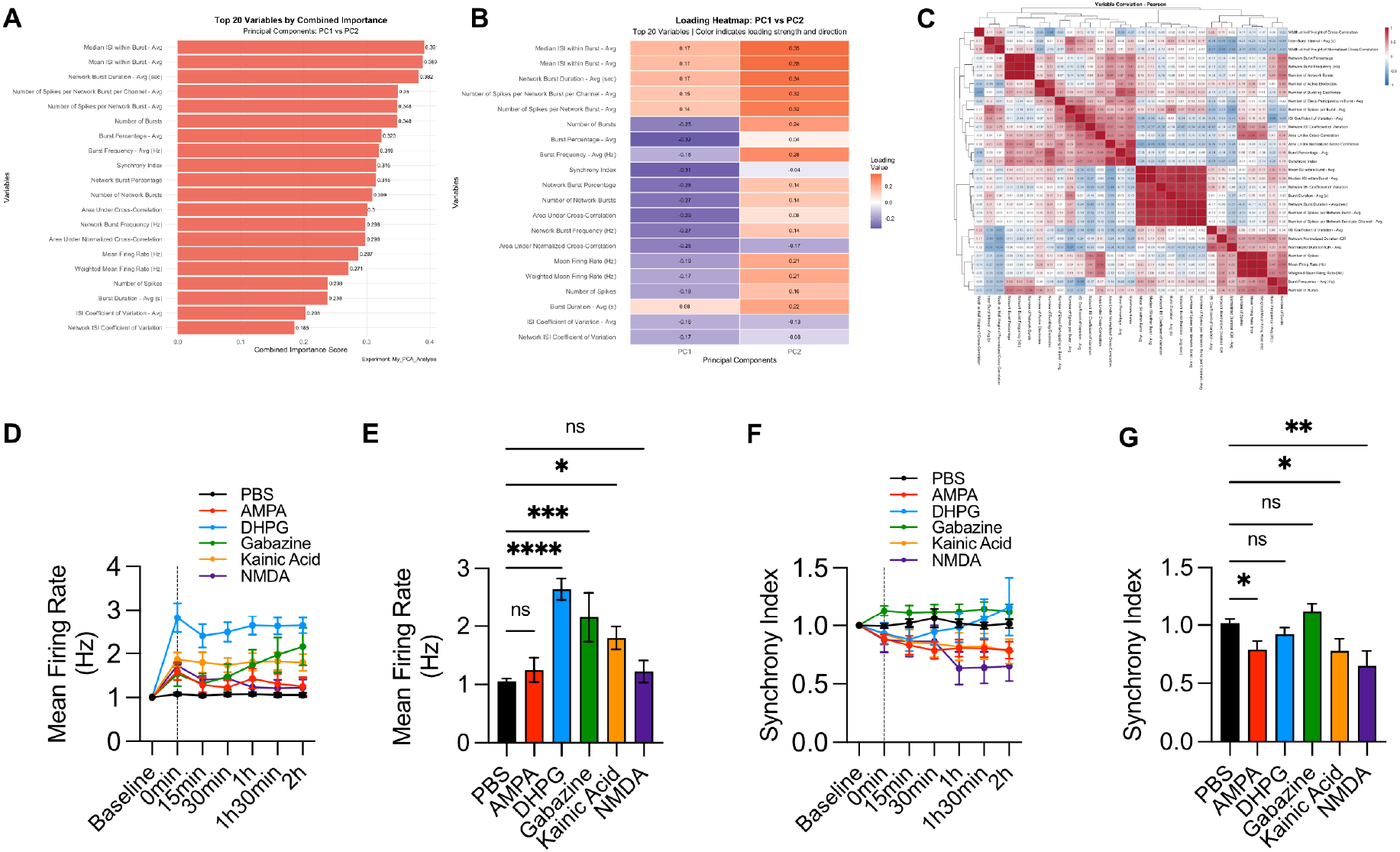
related to Figure 1. **(A)** Relative contributions of top 20 variables to Principal Component 1 and Principal Component 2 using NOVA for PCA analysis of neuronal agonists. **(B)** Loading heatmap of PC1 vs PC2 of the top 20 variables. Loading color indicates loading strength and direction, using NOVA. **(C)** Correlation heatmap illustrating the variables that vary together with agonists treatment, using NOVA. **(D)** Synchrony index normalized to baseline after treatment with neuronal agonists. **(E)** Synchrony index normalized to baseline at T=2h after treatment (one-way ANOVA with Dunnett’s multiple comparisons). **(F)** Mean Firing Rate (Hz) normalized to baseline after treatment with neuronal agonists. **(G)** Mean Firing Rate (Hz) normalized to baseline at T=2h after treatment (one-way ANOVA with Dunnett’s multiple comparisons).

**Supplemental Figure 2:**
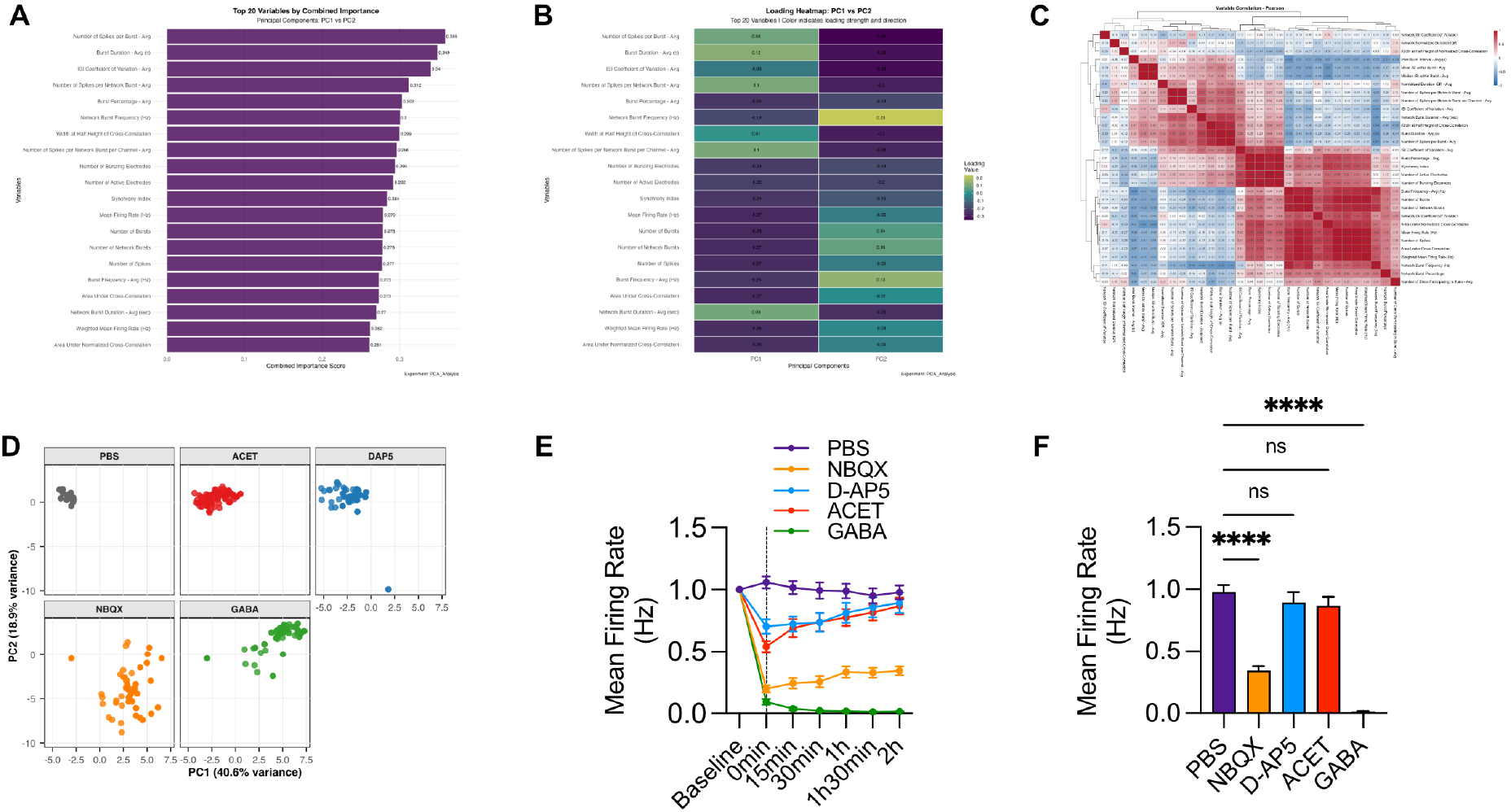
related to Figure 2. **(A)** NOVA representation: relative contributions of top 20 variables to Principal Component 1 and Principal Component 2 for the neuronal antagonists PCA analysis. **(B)** NOVA representation of loading heatmap of PC1 vs PC2 of the top 20 variables. Loading color indicates loading strength and direction. **(C)** NOVA representation of correlation heatmap illustrating the variables that vary together with antagonist treatment. **(D)** NOVA PCA plots split by treatment. Each dot represents a well. **(E)** Mean Firing Rate (Hz) normalized to baseline after treatment with neuronal antagonists. **(F)** Mean Firing Rate (Hz) normalized to baseline at T=2h after treatment (one-way ANOVA with Dunnett’s multiple comparisons).

**Supplemental Table 1.**

| **Metrics exported by Axis Navigator (n=44)** | **Metrics analyzed by NOVA (n=31)** |
| --- | --- |
| Number of Spikes | Number of Spikes |
| Mean Firing Rate (Hz) | Mean Firing Rate (Hz) |
| Number of Active Electrodes | Number of Active Electrodes |
| Weighted Mean Firing Rate (Hz) | Weighted Mean Firing Rate (Hz) |
| ISI Coefficient of Variation - Avg | ISI Coefficient of Variation - Avg |
| Number of Bursts | Number of Bursts |
| Number of Bursting Electrodes | Number of Bursting Electrodes |
| Burst Duration - Avg (s) | Burst Duration - Avg (s) |
| Burst Duration - Std (s) |  |
| Number of Spikes per Burst - Avg | Number of Spikes per Burst - Avg |
| Number of Spikes per Burst - Std |  |
| Mean ISI within Burst - Avg | Mean ISI within Burst - Avg |
| Mean ISI within Burst - Std |  |
| Median ISI within Burst - Avg | Median ISI within Burst - Avg |
| Median ISI within Burst - Std |  |
| Inter-Burst Interval - Avg (s) | Inter-Burst Interval - Avg (s) |
| Inter-Burst Interval - Std (s) |  |
| Burst Frequency - Avg (Hz) | Burst Frequency - Avg (Hz) |
| Burst Frequency - Std (Hz) |  |
| Normalized Duration IQR - Avg | Normalized Duration IQR - Avg |
| Normalized Duration IQR - Std |  |
| IBI Coefficient of Variation - Avg | IBI Coefficient of Variation - Avg |
| IBI Coefficient of Variation - Std |  |
| Burst Percentage - Avg | Burst Percentage - Avg |
| Burst Percentage - Std |  |
| Number of Network Bursts | Number of Network Bursts |
| Network Burst Frequency (Hz) | Network Burst Frequency (Hz) |
| Network Burst Duration - Avg (sec) | Network Burst Duration - Avg (sec) |
| Network Burst Duration - Std (sec) |  |
| Number of Spikes per Network Burst - Avg | Number of Spikes per Network Burst - Avg |
| Number of Spikes per Network Burst - Std |  |
| Number of Elecs Participating in Burst - Avg | Number of Elecs Participating in Burst - Avg |
| Number of Elecs Participating in Burst - Std |  |
| Number of Spikes per Network Burst per Channel - Avg | Number of Spikes per Network Burst per Channel - Avg |
| Number of Spikes per Network Burst per Channel - Std |  |
| Network Burst Percentage | Network Burst Percentage |
| Network IBI Coefficient of Variation | Network IBI Coefficient of Variation |
| Network ISI Coefficient of Variation | Network ISI Coefficient of Variation |
| Network Normalized Duration IQR | Network Normalized Duration IQR |
| Area Under Normalized Cross-Correlation | Area Under Normalized Cross-Correlation |
| Area Under Cross-Correlation | Area Under Cross-Correlation |
| Width at Half Height of Normalized Cross-Correlation | Width at Half Height of Normalized Cross-Correlation |
| Width at Half Height of Cross-Correlation | Width at Half Height of Cross-Correlation |
| Synchrony Index | Synchrony Index |

